# Identification and Analysis of Novel RNA Editing Sites in Neurodegenerative Diseases Using Machine Learning Approaches

**DOI:** 10.64898/2026.04.09.716726

**Authors:** Shabistan Jabin, Elamathi Natarajan

**Affiliations:** Biotecnika Info Labs PVT Ltd

**Keywords:** Control, Alzheimer’s disease (AD), RNA-seq, novel, ADAR, AEI

## Abstract

**Background:** RNA editing is a post-transcriptional modification that alters the sequence of an RNA transcript. Two types of RNA editing were found in mammals, involving the enzymatic deamination of either adenosine to inosine (A-to-I) or cytidine to uridine (C-to-U) nucleotides in RNA. A-to-I, which is the most common form of RNA editing, is mediated by the ADAR (adenosine deaminases acting on RNA) family of enzymes, ADAR1, ADAR2, and ADAR3. The editing event alters the hydrogen bond pairing of nucleobases, and the editing site will be recorded as guanosine rather than the original adenosine. Indeed, RNA editing deregulation has been linked to several nervous and neurodegenerative diseases. In this project work is done on Alzheimer’s disease (AD) and the samples are from anterior cingulate cortex of human brain tissue. AD is the main dementia in the world and a neurodegenerative condition prevalent in the elderly.

**Methodology:** A total of 20 raw RNA-sequencing data samples containing 10 controls and 10 Alzheimer’s disease (AD) cases were collected from NCBI using SRA Toolkit. Quality assessment was performed using FastQC and processed using Trimmomatic. Alignment was done using STAR RNA-seq aligner. RNA editing detection was performed using REDItools, detected sites were subsequently annotated against the REDIportal database. The resulting control-specific and disease-specific novel editing sites were merged into a single dataset containing exclusively novel, group-specific A-to-I editing events. This merged dataset was subsequently used for downstream feature extraction and machine learning analysis. Probability-based filtering was done to extract high-confidence disease associated sites and their gene list was used for computational level biological validation, pathway and functional enrichment analysis as well as overlap with known AD loci.

**Results:** Random Forest showed the highest accuracy score (0.804) and ROC-AUC score (0.854). Most important features that differentiated control and diseased novel sites in random forest were coverage (∼0.35), editing level (∼0.33) and GC content (∼0.15). The AEI mean values is higher in both male and female diseased cases (∼0.48-0.50) but less in male and female control cases (∼0.14-0.21). The mean values of ADAR1_CPM higher in control cases (123.65-143.30) and is less in diseased cases (88.35-97.93), ADAR2_CPM is almost equal in all cases (∼3.7-4.7) and ADAR3_CPM is very less in all the cases (∼0-0.02). Most candidate editing site were present in exon (∼62-67 %) CDS regions (∼17–21%) and relatively smaller fraction of gene (∼15-16 %). Editing alterations preferentially affect molecular systems governing synaptic structure, neurotransmission, and central nervous system integrity. In the main set -of the 2576 high-confidence genes identified, 33 overlapped with AD GWAS loci. In the core set -of the 1367 high-confidence genes identified, 11 overlapped with AD GWAS loci.

**Conclusion:** Feature like coverage, editing level and GC content contributed most. Alu sites are negligible as compared to non-alu sites but the AEI mean values are higher in diseased cases than in control cases. The mean values of ADAR1_CPM are higher than ADAR2_CPM and ADAR3_CPM.Sex does not play a major factor. High-confidence disease-associated RNA editing sites are strongly biased toward transcript-centric regions, particularly exons, with a notable subset affecting coding sequences. Importantly, enrichment of neurodegeneration-associated pathways and cognition-related human phenotypes further supports the disease relevance of these gene networks. RNA editing events in Alzheimer’s cortex may represent a regulatory mechanism largely independent of inherited genetic susceptibility loci.

## INTRODUCTION

RNA editing is a post-transcriptional modification that alters the sequence of an RNA transcript. It was first discovered by Benne et al. in a mitochondrion-encoded mRNA in *Trypanosoma brucei*.[1] Wagner et al. showed that RNA editing also occurs in mammalian cells. [2]

In fact, two types of RNA editing were found in mammals, involving the enzymatic deamination of either adenosine to inosine (A-to-I) or cytidine to uridine (C-to-U) nucleotides in RNA. [2–4]

Inosine is commonly interpreted as guanosine by translation and splicing machineries other than sequencing enzymes. As a consequence, A-to-I modifications can alter codon identity and increase transcriptome as well as proteome diversity.[5]

A-to-I, which is the most common form of RNA editing, is mediated by the ADAR (adenosine deaminases acting on RNA) family of enzymes, ADAR1, ADAR2, and ADAR3. [6,7] ADAR1 and ADAR2 are catalytically active. ADAR1 is ubiquitously expressed while ADAR2 expression is very low in some tissues. ADAR3 has no proven catalytic activity, and its expression is limited to the brain.[8] ADAR3 has been shown to inhibit ADAR1-induced RNA editing.[9] ADAR1 and ADAR2, but not ADAR3, deaminate adenosines to inosines, which are interpreted as guanosines by the translational and splicing machinery.[6] ADARs likely have non-editing-related functions too, but these are not discussed here.

In humans, three ADAR genes have been characterized: ADAR1 and ADAR2 encode for active enzymes and are expressed in most tissues and ADAR3, which does not seem to encode for a functional protein and is expressed exclusively in the central nervous system. [10]

Adenosine-to-inosine (A-to-I) RNA editing is a post-transcriptional processing event involved in diversifying the transcriptome and is responsible for various biological processes. A-to-I editing has now been identified as a reliable, differential biomarker in a number of neurological disorders. [11]

Indeed, RNA editing deregulation has been linked to several nervous and neurodegenerative diseases such as epilepsy, schizophrenia, major depression and amyotrophic lateral sclerosis [12,13].

The editing event alters the hydrogen bond pairing of nucleobases, and the editing site will be recorded as guanosine rather than the original adenosine. Moreover, A-to-I editing events are related to numerous critical biological processes, such as amino acid alterations [6], RNA splicing [6,14], nuclear retention [15], RNA interference [6,16,17] and innate cellular immunity [6,18,19]. RNA editing events may affect RNA localization, structure, stability, and transcript processing of both coding and noncoding RNAs (ncRNAs).[20] Editing is crucial for cell and tissue homeostasis and a variety of human diseases, including both cardiovascular and neurological disorders, have been linked to its deregulation. [21-25]

In present computational methods, A-to-G fake positive signals possibly result from sequencing errors, SNPs, somatic mutations, unfavourable amplification of pseudogenes, PCR errors and spurious chemical alterations in RNA. [26]

Low-expression transcripts and low-editing level sites may be ignored after rigorous bioinformatics screening from low coverage RNA-seq data. Therefore, extensive computational screening is necessary to predict low-editing rate A-to-I sites.[11]

In this project work is done on Alzheimer’s disease (AD) and the samples are from anterior cingulate cortex of human brain tissue.

Alzheimer’s disease (AD) is often characterized by cognitive impairment,[27] which may be accompanied by other symptoms, including sleep disturbances,[28] mood disorders,[29] autonomic dysfunctions,[30] specific sensory disorders (olfaction, hearing, vision),[31] and others.

AD is the main dementia in the world and a neurodegenerative condition prevalent in the elderly.[32,33] Despite the research progress since Alois Alzheimer’s inaugural description of the disease in1906,[34] the consensus amongst researchers is that AD pathology influences mostly in the cortex and select subcortical brain regions, including the frontal, temporal, and parietal lobes, portions of the cingulate gyrus, the hippocampus, and certain brainstem nuclei.[35]

The core pathogenesis of AD involves amyloid-β (Aβ) deposition, hyperphosphorylated tau (p-Tau), neuroinflammation, oxidative stress, [36,37] and others. Numerous neurons undergo degeneration and cell death in AD. [38]

## METHODOLOGY

### Human RNA-sequencing Data

A total of 20 raw RNA-sequencing data samples were collected from NCBI [39] and FASTQ files were downloaded via Sequence Read Archive (SRA) using SRA Toolkit (v 3.2.1) or directly from European Nucleotide Archive (ENA)(www.ebi.ac.uk).

The dataset comprised Illumina paired-end RNA-Seq libraries generated from human (*Homo sapiens*) anterior cingulate cortex brain tissue samples collected from individuals aged 55–90 years. Each sample was sequenced to a depth of approximately 50 million (M) 2 × 100 bp paired-end reads across two sequencing lanes.

The 20 samples included 10 neurologically normal controls and 10 Alzheimer’s disease (AD) cases further classified into 4 groups-control male, control female, diseased male and diseased female with 5 samples per group.

### Preprocessing Raw Data

#### Quality Control and Read Trimming

High-throughput sequencing quality assessment was performed using FastQC (v0.12.1). Raw paired-end FASTQ files were processed using Trimmomatic (v0.39) in paired-end mode for adapter removal and quality filtering. Illumina TruSeq adapter sequences were removed using the ILLUMINACLIP module (TruSeq3-PE.fa:2:30:10). Low-quality bases with Phred scores below 5 were trimmed from the beginning and end of reads (LEADING:5; TRAILING:5). A sliding window approach (SLIDINGWINDOW:4:20) was applied to remove regions where the average quality within a 4-base window fell below a Phred score of 20. Reads shorter than 50 bp after trimming were discarded (MINLEN:50).

Both paired and unpaired reads were retained for downstream analysis. Trimming was performed with multi-threading enabled where applicable.

#### Read Alignment to the Reference Genome

Only high-quality paired reads were retained for downstream analysis and aligned to the human reference genome using STAR RNA-seq aligner (Galaxy Version 2.7.11b) implemented on the Galaxy platform (usegalaxy.org).

Reads were mapped to the human reference genome GRCh38/hg38 (b38) using the built-in genome index. Splice junction annotation was provided using the GENCODE v45 gene annotation (gencode.v45.annotation.gtf).

Alignment was performed in paired-end mode (forward and reverse reads provided as individual datasets). Two-pass mapping was enabled to improve novel splice junction detection. Chimeric alignments were not reported. Standard alignment tags (NH, HI, AS, nM, NM, MD, jM, jI, MC, XS) were included in the BAM output. Unmapped reads were excluded from the final BAM files. Default alignment and seed parameters were used. Strand-specific genome coverage files were generated in bedGraph format and normalized to reads per million mapped reads (RPM).

The resulting alignment log files were subjected to quality assessment using FastQC on individual samples, followed by aggregate quality summarization using MultiQC for group-wise evaluation and the BAM files were used as input for group-wise gene-level quantification using featureCounts on the Galaxy platform (usegalaxy.org).

#### Identification of A-to-I RNA Editing Sites

Individual BAM alignment files were downloaded from the Galaxy platform and subjected to RNA editing analysis using the REDITools v2.0 pipeline. A-to-I RNA editing sites were identified following the standard filtering strategy implemented in REDItools. [40]

REDItools is the first published software for genome-wide RNA ed iting site detection and provides both A-to-I and C-to-U variant detection. It is suitable for both RNA-seq and DNA-seq data from the same sample/individual or RNA-seq data alone and uses pre-aligned reads in Binary Alignment/Map (BAM) format as input. It performs de novo and known editing site detection and utilizes a wide range of filters and quality control checks for the identification of false positives, especially near intronic splice sites, read-ends and homopolymeric regions. REDItools provides additional scripts for post-processing output tables.[41]

To retain primary alignments mapped to canonical chromosomes, BAM files were filtered using samtools to include reads mapped to chromosomes 1–22, X, and Y. The filtered BAM files were indexed prior to downstream analysis.

RNA editing detection was performed using the human reference genome GRCh38 primary assembly. REDItools was executed with the following parameters: minimum read depth ≥ 20 (-l 20), minimum base quality ≥ 30 (-q 30), minimum supporting reads ≥ 2 (-s 2), removal of duplicate reads (-u), exclusion of multi-mapped reads (-e), and multi-threading enabled (-t 4).

Candidate A-to-I editing events were defined as A-to-G mismatches relative to the reference genome.

The resulting REDItools files were classified group wise as control male, control female, diseased male and diseased female with 5 files per group.

A-to-I RNA editing sites were initially identified from the STAR-aligned BAM files using REDItools. The detected sites were subsequently annotated against the REDIportal database using TABLE1_hg38_v3 to determine previously reported editing events.[42]

Editing sites were classified into two categories:

- **Known A-to-I sites**: Sites present in the REDIportal database.
- **Novel A-to-I sites**: Sites not reported in REDIportal.

This classification enabled downstream comparative analyses between known and potentially novel RNA editing events across experimental groups.

#### Identification of Group-Specific Novel Editing Sites

Further filtering was performed using the subset of sites classified as novel (i.e., not reported in REDIportal). To identify group-specific editing events, novel sites detected in control samples were compared against those detected in Alzheimer’s disease (AD) samples.

Control-specific novel sites were defined as editing sites present in control samples but absent in both REDIportal and all AD REDItools outputs. Conversely, disease-specific novel sites were defined as sites present in AD samples but absent in both REDIportal and all control REDItools outputs.

The resulting control-specific and disease-specific novel editing sites were merged into a single dataset containing exclusively novel, group-specific A-to-I editing events. This merged dataset was subsequently used for downstream feature extraction and machine learning analysis.

Sites were required to have a minimum Q30 coverage of ≥10 reads and an editing frequency of ≥10% (0.1). Sites lacking valid editing frequency values were excluded. Most of it is done in REDItools.

REDItools output files were filtered to retain only canonical A-to-I editing events, defined as A→G substitutions on the sense strand and T→C substitutions on the antisense strand. The reference base column containing A and T bases is added in the final table.

These criteria ensured the selection of high-confidence and biologically relevant RNA editing events

### Feature Engineering

#### Addition of Sample Metadata and Functional Annotation

Condition and sex columns were added to the final table. Gene information (gene name and Ensembl IDs) and genomic region (exon, intron, UTR, intergenic, etc.) were added after functional annotation using GENCODE v45 (gencode.v45.annotation.gtf). Each site was labelled according to its genomic location (exon, intron, UTR, or intergenic).

#### Alu and non-alu repeats

Most RNA editing occurs in Alu repeats. Editing sites overlapping Alu repeats were identified using RepeatMasker annotations (hg38). RepeatMasker(hg38) was downloaded from UCSC [43]and only Alu elements were extracted. Alu flag column (0/1) is added denoting alu as 1 and non-alu as 0.

#### ±5 bp sequence and derivation of numeric features

Sequences were extracted from reference genome (hg38.fa) keeping the window length of 11 bp (5 bp upstream + edited base + 5 bp downstream) and compute sequence features.

We derived interpretable numeric features viz GC content, motif AG (0/1), motif AA (0/1), motif TT (0/1).

#### ADAR Counts Per Million

Sample-level expression of RNA editing enzymes ADAR1, ADAR2, and ADAR3 was quantified using featureCounts output files. Raw counts were normalized to counts-per-million (CPM) and included as covariates in downstream modelling to account for inter-sample variability in RNA editing activity.

Raw gene counts along with Ensembl IDs for ADAR genes were extracted from the featureCounts output files. The total number of assigned reads per sample was obtained from the corresponding featureCounts summary files.

CPM (Counts Per Million):

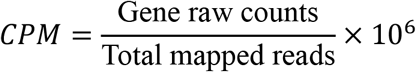

Since ADAR acts globally rather than in a site-specific manner, expression levels were calculated per sample, and the mean value was subsequently computed for each group.

#### AEI (Alu Editing Index)

Global Alu Editing Index (AEI) was computed using editing level and Alu flag columns per condition and sex as the mean A-to-I editing level across all Alu-associated sites. AEI was used as a sample-level covariate to contextualize novel RNA editing events in neurodegenerative disease.

AEI per sample group:

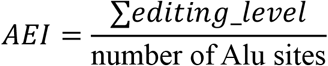

The mean non-Alu editing level was calculated group-wise to represent background editing occurring outside Alu regions.

### Machine Learning

A data table containing relevant features was created for machine learning analysis. Since the dataset was supervised, supervised learning models were applied. Three models—Logistic Regression, Random Forest, and XGBoost—were designed and saved.

Only novel editing sites were included in the dataset, and the models were trained to classify control-specific novel sites and disease-specific novel sites. For each model, the classification report, accuracy score, and ROC-AUC score were calculated.

Among the three models, the Random Forest model showed the highest accuracy; therefore, subsequent analyses were performed using this model.

#### Probability-based filtering

High-confidence disease-predicted novel editing sites were selected from the full dataset using probability-based filtering. A threshold of ≥0.90 was applied to define the main set, while a more stringent threshold of ≥0.95 was used to define the core set. These represent the final predicted disease-associated editing sites.

The core set (≥0.95) is a strict subset of the main set (≥0.90), containing fewer but more confidently predicted sites.

Region-wise counts (exon, CDS, and gene) were extracted from both sets for comparative analysis. Gene lists were also obtained from each set for downstream functional and pathway analyses of the candidate editing sites.

### Biological validation (computational level)

#### Pathway and Functional Enrichment Analysis

Functional enrichment analysis was performed using g:Profiler GOSt on the Galaxy platform (usegalaxy.org). Gene lists derived from both the main set (probability ≥0.90) and the core set (probability ≥0.95) were analysed.

Functional enrichment analysis was performed using g:Profiler GOSt on the Galaxy platform (usegalaxy.org). Gene lists from both the sets (main set and core set) were analysed for enrichment against Gene Ontology (GO) categories, including Biological Process (BP), Molecular Function (MF), and Cellular Component (CC), as well as biological pathway databases such as KEGG, Reactome, and WikiPathways. Disease/phenotype database: Human Phenotype Ontology (HP) terms were also assessed.

Only statistically significant terms were retained using the default false discovery rate (FDR) threshold of 0.05. All other parameters were kept at default settings.

This analysis provided functional annotation, pathway enrichment, and phenotype association of the candidate genes, serving as computational biological validation of the machine learning– selected disease-associated editing sites.

#### Overlap with Known Disease Loci

To assess genetic consistency, overlap analysis was performed with known Alzheimer’s disease (AD) loci. Since the samples were derived from Alzheimer’s anterior cingulate cortex brain tissue, enrichment was specifically evaluated against Alzheimer’s GWAS findings.

The GWAS Catalog associations file was downloaded, and Alzheimer’s-associated genes were extracted using a genome-wide significance threshold of p ≤ 5 × 10^−8^.

Overlap was computed separately for:

- Core set genes (probability ≥ 0.95)
- Main set genes (probability ≥ 0.90)

Statistical significance of the overlap was assessed using a hypergeometric test, which evaluates whether the observed overlap is greater than expected by chance.

Parameters:

- N = total protein-coding genes (∼20000)
- K = number of GWAS genes
- n = number of main/core genes
- k = observed overlap Expected Random Overlap

Formula:

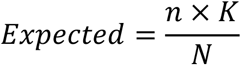

## RESULTS AND DISCUSSION

In ML, among the three models—namely Logistic Regression, Random Forest, and XGBoost— Random Forest showed the highest accuracy score (0.804) and ROC-AUC score (0.854).

**graph.1.**
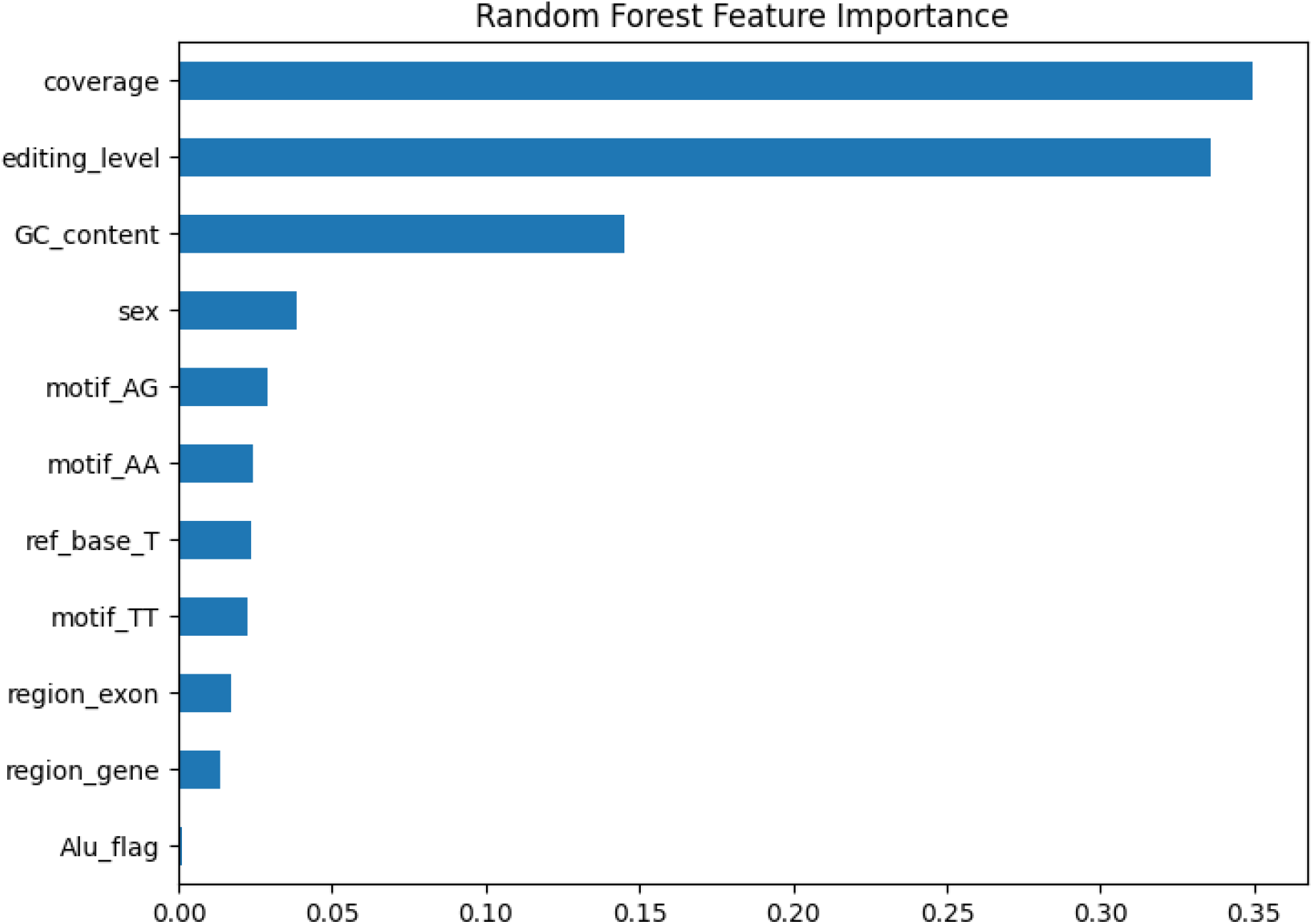
Random Forest Feature Importance. The graph.1 shows most important features that differentiated control and diseased novel sites in random forest are: Coverage (∼0.35) Editing level (∼0.33) GC content (∼0.15) Everything else contributes much less.

**graph.2.**
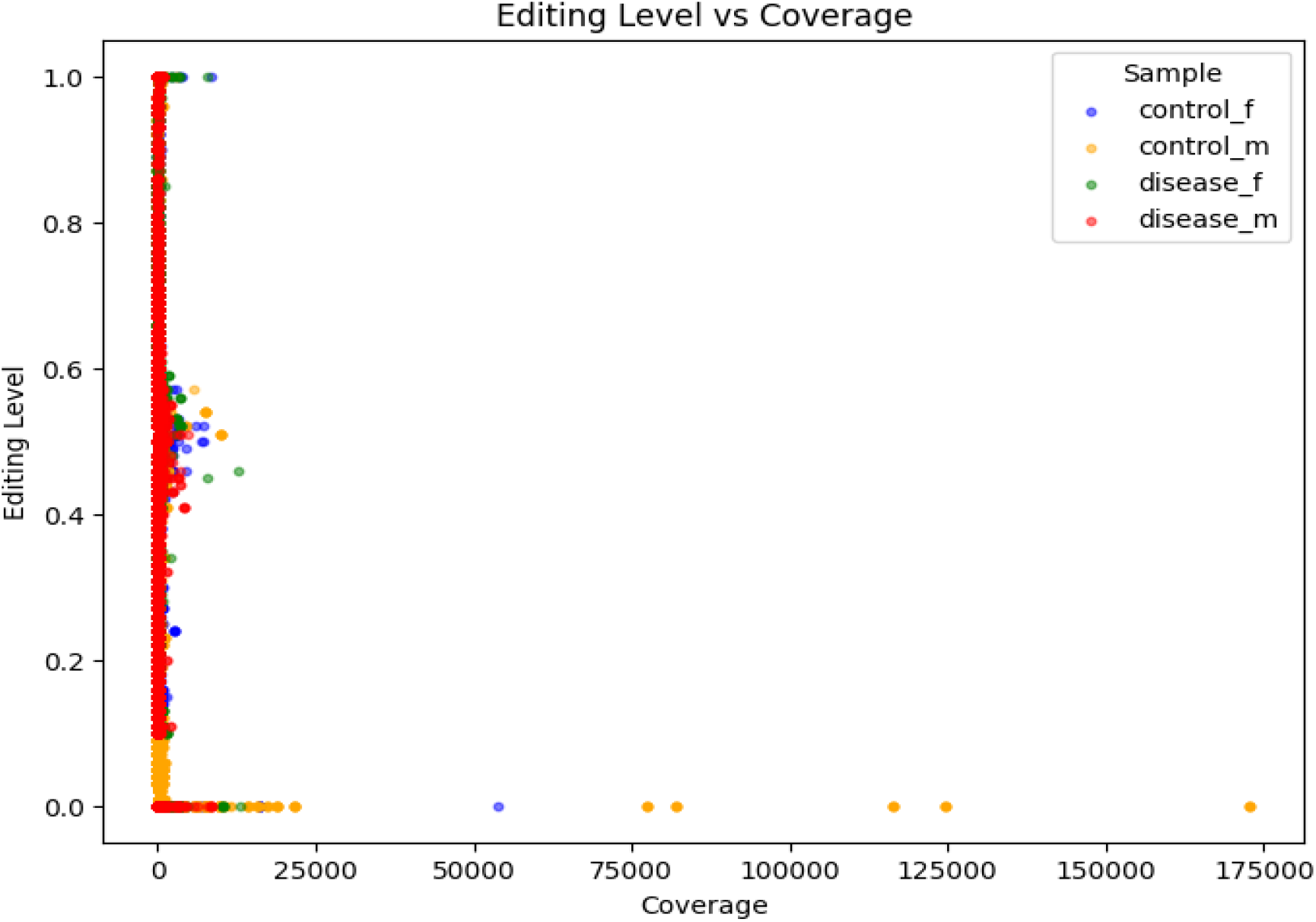
Editing Level vs Coverage. The graph.2 shows relation between editing level and coverage for control novel and diseased novel editing sites. Most of the sites that occupy editing level (∼0.5-1.0) are of very low coverage (∼10-12,000). Higher coverage shows 0 editing level. A high number of disease male and disease female sites cluster at significant editing level (∼0.5-1.0).

**graph.3.**
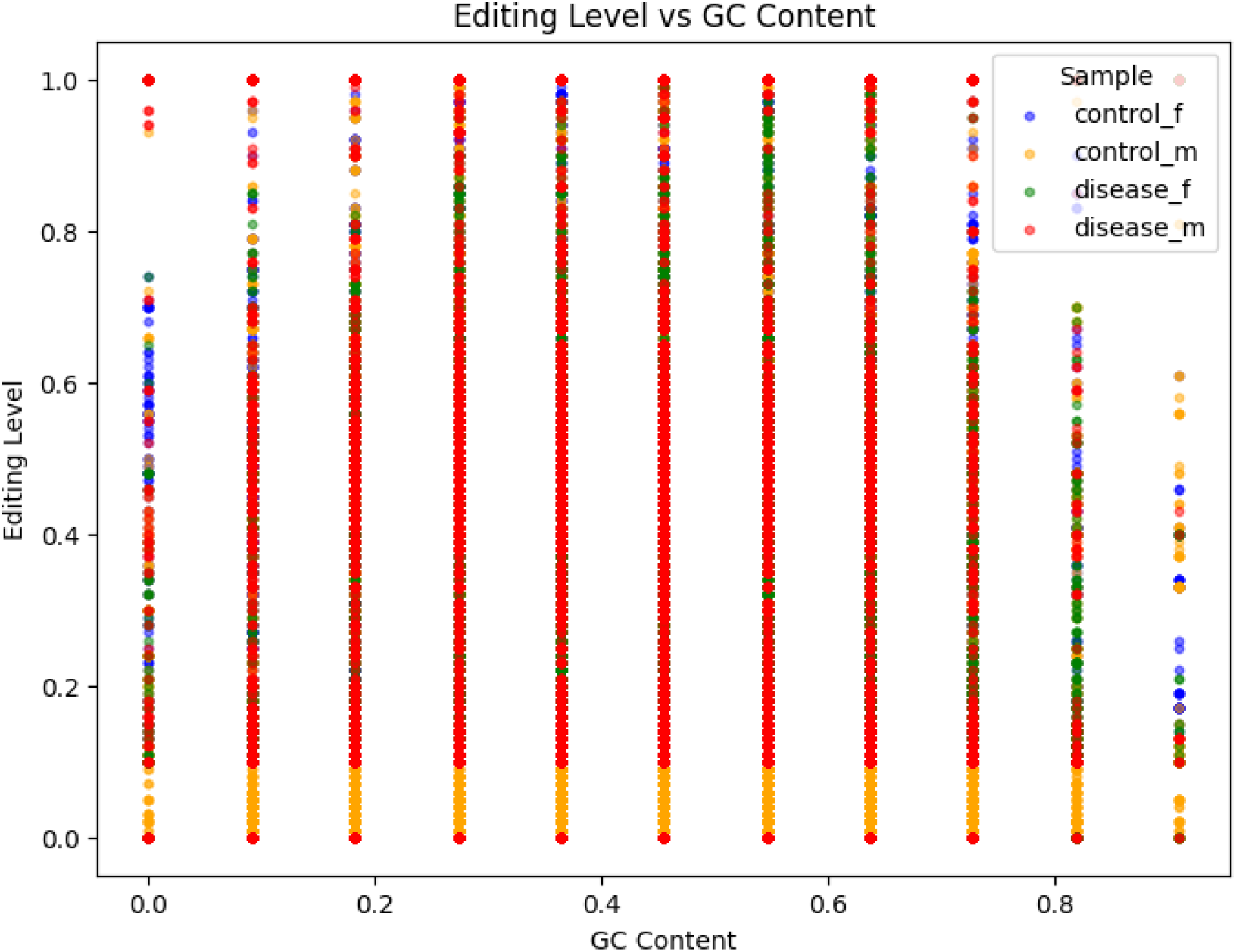
Editing Level vs GC Content. The graph.3 shows relation between editing level and GC content for control novel and diseased novel editing sites. A high number of disease male and moderate number disease female sites cluster at significant editing level (∼0.5-1.0) with the GC content ranging from 0 to 1.

**graph.4.**
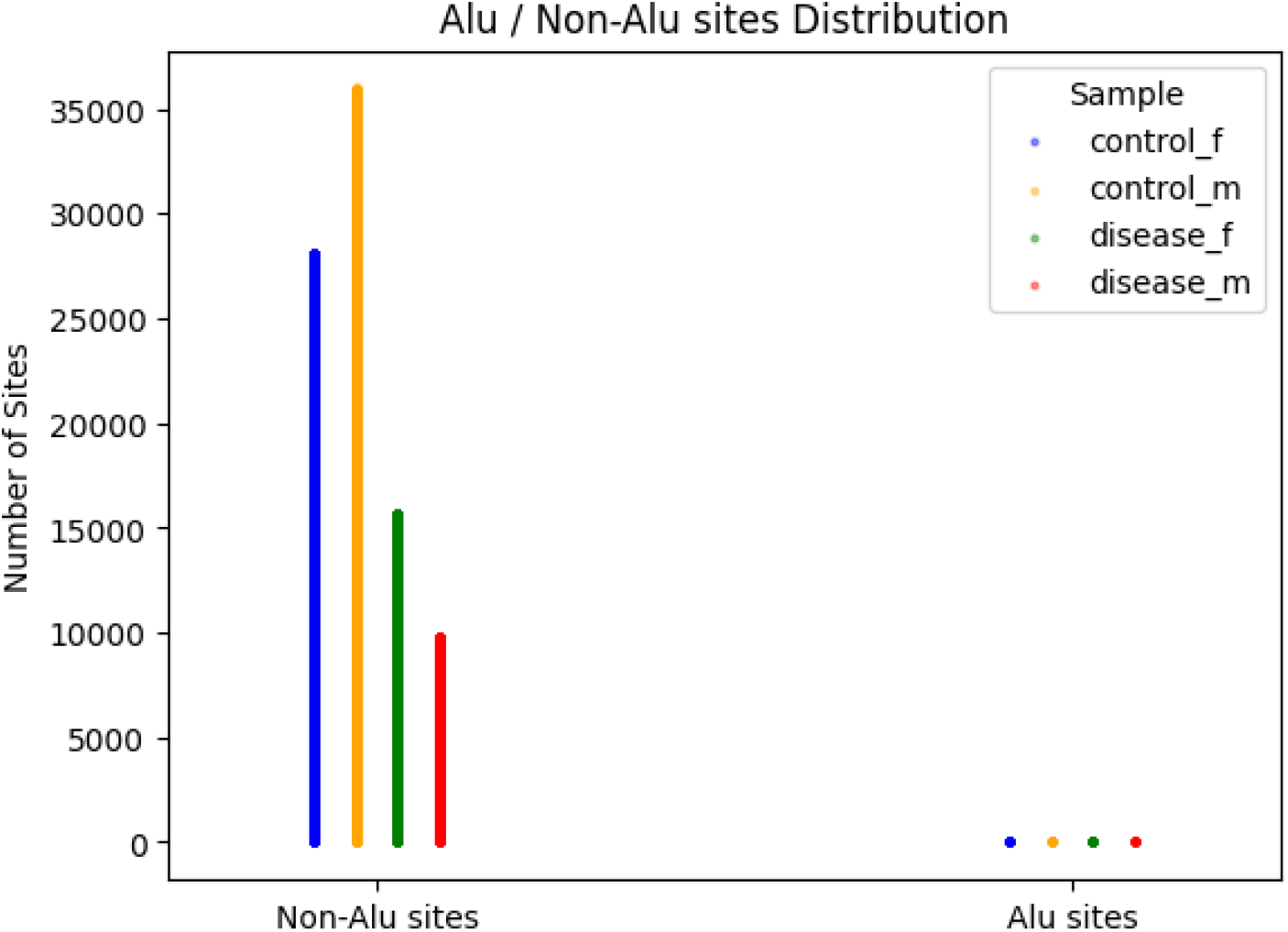
Alu/Non-Alu Sites Distribution. The graph.4 shows that the number of non-alu sites are higher in comparison to the alu sites and the number of non-alu sites are higher in control cases (∼27,000-36,000) than in diseased cases(∼9,000-15,000).

**graph.5.**
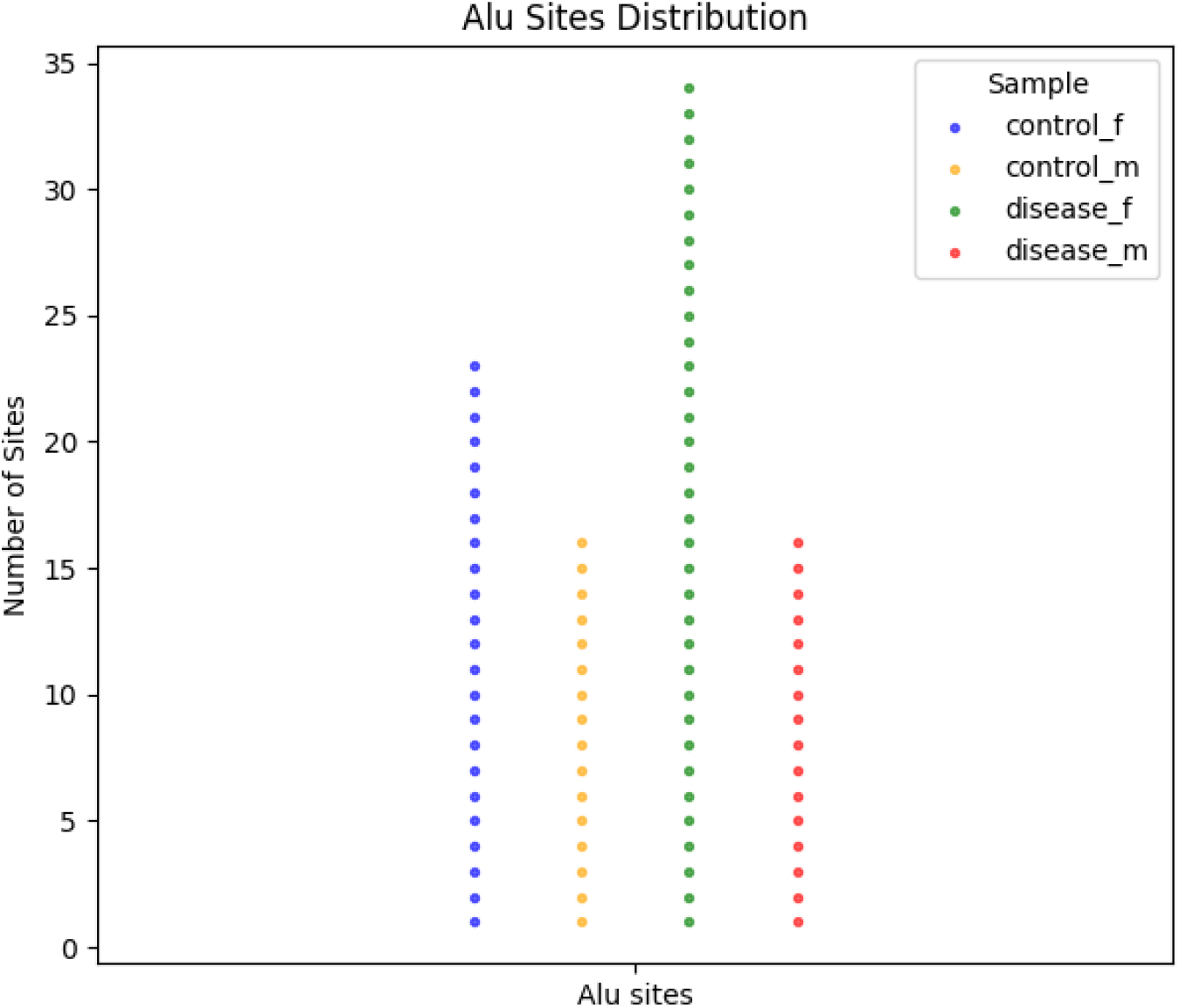
Alu Sites Distribution. The graph.5 shows the number of alu-sites are higher in control female (23) and diseased female (34) cases in comparison to control male (16) and diseased male (16) cases.

**Table 1.**
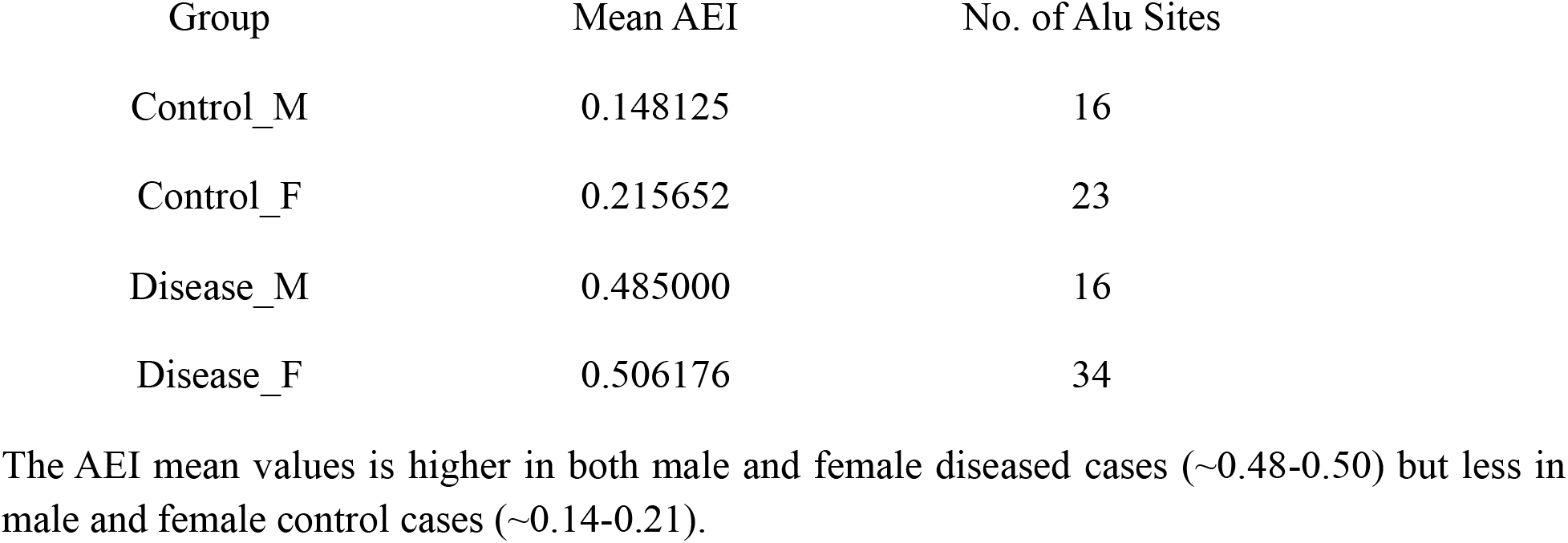
AEI Mean Values Group-wise.

| Group | Mean AEI | No. of Alu Sites |
| --- | --- | --- |
| Control_M | 0.148125 | 16 |
| Control_F | 0.215652 | 23 |
| Disease_M | 0.485000 | 16 |
| Disease_F | 0.506176 | 34 |

The AEI mean values is higher in both male and female diseased cases (~0.48-0.50) but less in male and female control cases (~0.14-0.21).

**Table 2.** ADAR Mean Values Group-wise.

| Group | ADAR1_CPM | ADAR2_CPM | ADAR3_CPM |
| --- | --- | --- | --- |
| Control_M | 143.30906 | 4.681654 | 0.0206042 |
| Control_F | 123.657682 | 3.885048 | 0.00858442 |
| Disease_M | 97.9317074 | 3.789548 | 0 |
| Disease_F | 88.3565932 | 4.306736 | 0.01033818 |

The mean values of ADAR1\_CPM higher in control cases (123.65-143.30) and is less in diseased cases (88.35-97.93), ADAR2\_CPM is almost equal in all cases (~3.7-4.7) and ADAR3\_CPM is very less in all the cases (~0-0.02).

In the feature extraction dataset control novel sites were 64,076 and disease novel sites were 25,590. After probability threshold filtering the number of high-confidence disease sites in main set was 3201 (12.5%) with 2576 genes and number of high-confidence disease sites in core set was 1558 (6.1%) with 1367 genes. The core set is a complete subset of the main set.

**Table 3.** Genomic Distribution of High-Confidence Disease-Associated RNA Editing Sites.

**Main Set (0.90) — 3201 sites**
| Region | Count | % |
| --- | --- | --- |
| exon | 2014 | 62.90% |
| CDS | 663 | 20.70% |
| gene | 524 | 16.40% |
| total | 3201 |  |

**Core Set (0.95) — 1558 sites**
| Region | Count | % |
| --- | --- | --- |
| exon | 1043 | 67.00% |
| CDS | 272 | 17.50% |
| gene | 243 | 15.60% |
| total | 1558 |  |

To investigate the genomic localization of predicted disease-associated RNA editing sites, we examined their distribution across annotated regions (exon, CDS, and gene body) in both the main set (n = 3201) and the core set (n = 1558). In the main set, the majority of editing sites were located within exonic regions (62.9%), followed by coding sequences (CDS; 20.7%) and gene body regions (16.4%). A similar pattern was observed in the core set, where 67% of sites mapped to exons, 17.5% to CDS, and 15.6% to gene body regions.

Notably, the predominance of exonic localization (∼62-67 %) was maintained and slightly increased at the higher confidence threshold, indicating a stable and biologically consistent signal. The substantial proportion of sites within CDS regions (∼17–21%) suggests potential functional consequences at the protein level, including possible amino acid substitutions and modulation of protein activity. Meanwhile, the relatively smaller fraction of gene body-associated sites indicates that disease-associated editing events are preferentially enriched within mature transcript regions rather than broadly distributed across gene loci.

### Functional Enrichment Analysis of the Main Gene Set

Comprehensive functional enrichment analysis of the high-confidence main gene set revealed a highly structured neuronal signature across Gene Ontology (GO), KEGG, Reactome, and Human Phenotype (HP) databases. Across all annotation systems, enrichment converged on synaptic organization, intracellular signalling cascades, cytoskeletal remodelling, vesicle trafficking, and cognitive phenotypes, indicating that the gene set forms a coherent neurobiological module rather than a random collection of genes.

#### Gene Ontology (GO) Enrichment Analysis

##### GO: Biological Process (BP)

Biological Process enrichment demonstrated strong overrepresentation of neuronal developmental and regulatory programs. The most significantly enriched categories included regulation of signal transduction, small GTPase-mediated signalling, synapse organization, neuron differentiation, axon genesis, dendrite development, cytoskeleton organization, vesicle-mediated transport, and regulation of synaptic plasticity. Higher-order processes such as learning, memory, cognition, and behaviour were also significantly enriched.

The repeated identification of cytoskeletal organization and Rho GTPase–mediated signalling highlights structural plasticity as a central axis of the gene set. Cytoskeletal remodelling underlies dendritic spine dynamics, synaptic stabilization, and activity-dependent adaptation. Given that dendritic spine loss and structural instability are early features of Alzheimer’s disease pathology, the enrichment pattern suggests that RNA editing–associated genes are embedded within molecular systems governing neuronal structural integrity and plasticity.

Importantly, the clustering of genes within regulatory and signalling processes indicates biological structure rather than stochastic enrichment, supporting functional coherence of the main set.

##### GO: Molecular Function (MF)

Molecular Function categories were dominated by protein binding, small GTPase binding, kinase binding, cytoskeletal protein binding, scaffold protein activity, and signalling adaptor activity. These functions emphasize regulatory and interaction-based molecular roles rather than enzymatic housekeeping activities.

The prominence of binding and adaptor functions suggests that the gene set participates in signal integration and modulation of intracellular cascades. Such molecular functions are characteristic of proteins that coordinate dynamic signalling events and structural remodelling within synapses. This aligns closely with the BP and CC findings, reinforcing a model of coordinated synaptic regulation.

##### GO: Cellular Component (CC)

Cellular Component enrichment revealed predominant localization of the genes to synaptic and neuronal subcellular compartments. Highly enriched terms included synapse, post-synapse, pre-synapse, dendrite, axon, neuron projection, cytoskeleton, actin cytoskeleton, postsynaptic density, and membrane-associated vesicular compartments.

This localization pattern demonstrates that the gene set spans both presynaptic and postsynaptic machinery, suggesting involvement in multiple layers of synaptic function. Enrichment within postsynaptic density and vesicle-related compartments reinforces participation in neurotransmission and synaptic structural organization. The spatial specificity observed in CC categories further supports the interpretation that these genes are concentrated within neuronal communication hubs rather than general cellular compartments.

#### KEGG Pathway Enrichment

KEGG pathway analysis revealed significant enrichment in pathways central to neuronal signalling and plasticity. Enriched pathways included glutamatergic synapse, GABAergic synapse, long-term potentiation, calcium signalling pathway, MAPK signalling pathway, PI3K-Akt signalling pathway, axon guidance, and regulation of actin cytoskeleton.

These pathways are fundamental to activity-dependent synaptic modulation and memory encoding. Enrichment of long-term potentiation directly links the gene set to mechanisms underlying learning and memory processes.

Notably, several neurodegeneration-related pathways, including Alzheimer’s disease and Parkinson’s disease, were significantly enriched. Although such pathways often contain broadly expressed neuronal components, their enrichment within this synaptic and signalling framework suggests mechanistic convergence with disease-associated molecular cascades. Rather than isolated disease labels, these pathways appear integrated within the broader synaptic regulatory architecture identified across databases.

#### Reactome Pathway Enrichment

Reactome analysis provided higher-resolution insight into the functional network and demonstrated strong enrichment in neuronal system pathways, transmission across chemical synapses, postsynaptic signal transmission, vesicle-mediated transport, Rho GTPase signalling, cytoskeletal remodelling, protein phosphorylation, and ubiquitin-mediated proteolysis.

The consistent identification of Rho GTPase cycle and cytoskeleton reorganization pathways across GO, KEGG, and Reactome analyses underscores structural plasticity as a recurring and central theme. Additionally, enrichment in phosphorylation and ubiquitination pathways indicates tight post-translational regulatory control of synaptic proteins.

Disruption of proteostasis and protein turnover mechanisms is a hallmark of neurodegenerative disorders. The presence of ubiquitin-mediated proteolysis and protein modification pathways within this framework suggests that RNA editing–associated genes may intersect with protein homeostasis systems that influence synaptic stability and vulnerability.

#### Human Phenotype (HP) Enrichment

Human Phenotype enrichment analysis demonstrated significant overrepresentation of neurological and cognitive phenotypes, including abnormal cognition, memory impairment, abnormal synaptic transmission, intellectual disability–related phenotypes, behavioural abnormalities, and neurodevelopmental traits.

This phenotypic convergence provides an independent validation layer linking molecular enrichment results to clinically relevant neurological manifestations. The alignment between synaptic plasticity pathways and cognition-related phenotypes strengthens the biological interpretation by connecting molecular systems to functional outcomes at the organismal level.

Across all enrichment layers, the 0.90 main gene set demonstrates consistent cross-database convergence on synaptic regulation, neuronal development, cytoskeletal remodelling, and intracellular signalling systems. The enrichment pattern is not driven by broad neuronal annotations but instead reflects structured organization within synaptic transmission pathways, structural plasticity mechanisms, and regulatory signalling cascades.

The simultaneous enrichment of excitatory and inhibitory synaptic pathways, Rho GTPase– mediated cytoskeletal dynamics, vesicle trafficking systems, and proteostasis mechanisms suggests that RNA editing–associated genes are positioned at critical nodes of neuronal adaptability. The overlap with neurodegeneration-related pathways and cognitive phenotypes further supports disease relevance.

Collectively, these findings indicate that the main 0.90 gene set represents a biologically coherent synaptic regulatory network, linking RNA editing–associated genes to structural remodelling, neurotransmission balance, and higher-order cognitive processes. This convergence across GO, KEGG, Reactome, and Human Phenotype ontologies provides strong functional validation of the identified editing-associated gene landscape in brain tissue.

### Functional Enrichment Analysis of the Core Gene Set

#### Gene Ontology Enrichment

##### GO: Biological Process (BP)

Biological Process enrichment revealed a strong neurodevelopmental and synaptic regulatory signature. Highly significant terms included nervous system development, neurogenesis, neuron differentiation, axon genesis, dendrite morphogenesis, neuron projection development, and central nervous system development.

Synaptic processes were prominently enriched, including synapse organization, post synapse organization, regulation of trans-synaptic signalling, modulation of chemical synaptic transmission, and positive regulation of glutamate secretion and neurotransmission. The presence of learning, memory, and cognition terms further connects this gene set to higher-order neural function.

In addition, cytoskeletal remodelling processes such as actin filament organization, neuron projection morphogenesis, and regulation of actin-based processes were significantly enriched. Intracellular signalling and phosphorylation-related pathways were also overrepresented, indicating dynamic signal modulation capacity.

Together, these results demonstrate that the 0.95 core gene set is enriched in coordinated neurodevelopmental, synaptic, and signalling programs rather than broad regulatory categories.

##### GO: Molecular Function (MF)

Molecular Function enrichment highlighted cytoskeletal protein binding and actin binding, consistent with structural remodelling capacity. Kinase binding, protein kinase activity, and phosphotransferase activity were enriched, indicating phosphorylation-mediated signalling mechanisms.

A prominent signal emerged for GTP binding, GTPase activity, small GTPase binding, and GTPase regulator activity. Small GTPases regulate dendritic spine morphology, vesicle trafficking, and cytoskeletal organization, providing mechanistic integration with BP and CC enrichment results.

Collectively, GO-MF results indicate that stringent editing-associated genes participate in kinase-driven signalling and GTPase-regulated cytoskeletal remodelling central to synaptic plasticity.

##### GO: Cellular Component (CC)

Cellular Component analysis showed strong localization to synaptic microdomains. Enriched compartments included synapse, neuron-to-neuron synapse, post-synapse, postsynaptic density, pre-synapse, dendritic spine, and glutamatergic synapse. Notably, enrichment of the Schaffer collateral–CA1 synapse suggests involvement in hippocampal circuitry associated with learning and memory.

Neuronal morphology compartments such as axon, dendrite, neuron projection, growth cone, and somatodendritic compartment were significantly overrepresented. Cytoskeletal structures including actin cytoskeleton and microtubule cytoskeleton were also enriched.

These findings indicate precise structural localization within neuronal architecture, particularly within synaptic and dendritic compartments critical for plasticity.

#### KEGG Pathway Enrichment

KEGG pathway analysis of the stringent 0.95 core gene set revealed significant enrichment in synaptic transmission pathways, including GABAergic synapse (p ≈ 0.0196) and Glutamatergic synapse (p ≈ 0.0477). These represent the two principal neurotransmission systems in the central nervous system. Additional enrichment in the morphine addiction pathway further reflects synaptic signalling architecture rather than metabolic processes.

The selective enrichment of GABAergic and glutamatergic pathways indicates that the machine learning–filtered genes are concentrated within core excitatory and inhibitory neurotransmission systems. Given the central role of excitatory–inhibitory balance in neurodegeneration, this enrichment suggests direct functional involvement of high-confidence editing-associated genes in synaptic circuitry regulation rather than generic cellular processes.

#### Reactome Pathway Enrichment

Reactome analysis demonstrated significant clustering in Protein–protein interactions at synapses (p ≈ 0.016) and the Neuronal System pathway (p ≈ 0.022). These pathways capture the molecular interaction networks governing synaptic structure, receptor signalling, and neurotransmission.

The convergence of Reactome synaptic interaction networks with KEGG synaptic transmission pathways reinforces that the stringent gene set is embedded within neuronal communication systems. This level of pathway specificity is consistent with disease-relevant brain biology and supports the biological validity of the ML-selected editing sites.

#### Human Phenotype (HP) Enrichment

Human Phenotype analysis demonstrated strong enrichment for central nervous system–related phenotypes, including neurodevelopmental abnormality, neurodevelopmental delay, abnormal nervous system physiology, seizure, abnormal cognitive process, abnormal mental function, abnormal brain morphology, abnormal forebrain morphology, and CNS atrophy/degeneration. The p-values reached highly significant levels (as low as ∼5.47 × 10^−7^).

Motor tone phenotypes (hypotonia, hypertonia) were also enriched, consistent with secondary manifestations of central nervous system dysfunction in neurodevelopmental and neurodegenerative disorders. Thus, phenotypic convergence strongly supports the neurological relevance of the ML-selected gene set.

Across GO, KEGG, Reactome, and Human Phenotype analyses, the stringent 0.95 core gene set demonstrates robust cross-database convergence on synaptic biology. The genes localize to synaptic compartments and dendritic spines, participate in glutamatergic and GABAergic neurotransmission pathways, regulate cytoskeletal remodelling through small GTPase and kinase signalling, and are strongly associated with neurodevelopmental and cognitive phenotypes.

This convergence indicates a highly coherent neurodevelopmental and synaptic regulatory program rather than random enrichment. Even in the absence of a direct “Alzheimer disease” KEGG label, the enrichment of excitatory/inhibitory synaptic pathways, neuronal system interaction networks, and CNS degeneration phenotypes provides mechanistic consistency with neurodegenerative biology.

Taken together, these findings support the biological relevance of stringent ML-filtered RNA editing–associated genes and suggest that editing alterations preferentially affect molecular systems governing synaptic structure, neurotransmission, and central nervous system integrity.

### Expected Random Overlap

Number of unique GWAS genes: **267**

#### For MAIN

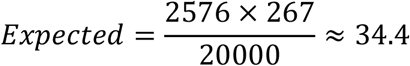

Observed = **33** (Almost exactly random expectation.)

Of the 2576 high-confidence genes identified, 33 overlapped with previously reported Alzheimer’s disease GWAS loci.

#### For CORE

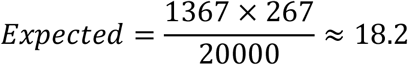

Observed = **11(**Lower than expected.)

Of the 1367 high-confidence genes identified, 11 overlapped with previously reported Alzheimer’s disease GWAS loci.

RNA editing–associated genes are not enriched for known Alzheimer’s GWAS risk loci. RNA editing alterations in Alzheimer’s cortex may represent a regulatory layer that is largely independent of inherited genetic risk loci.

Comparison with genome-wide significant Alzheimer’s disease GWAS loci revealed no significant enrichment among RNA editing–associated genes (main set: p = 0.63; core set: p = 0.98). The observed overlap did not exceed random expectation, suggesting that disease-associated RNA editing events in Alzheimer’s cortex may represent a regulatory mechanism largely independent of inherited genetic susceptibility loci.

## CONCLUSION

Feature like coverage, editing level and GC content contributed most.

Alu sites are negligible as compared to non-alu sites but the AEI mean values are higher in diseased cases than in control cases. The mean values of ADAR1_CPM are higher than ADAR2_CPM and ADAR3_CPM.

Sex does not play a major factor. Males and females are affected equally in Alzheimer’s disease.

The genomic localization of predicted disease-associated RNA editing sites findings demonstrate that high-confidence disease-associated RNA editing sites are strongly biased toward transcript-centric regions, particularly exons, with a notable subset affecting coding sequences. This distribution supports a functional role for RNA editing in modulating neuronal transcript architecture and potentially protein function in disease contexts.

Taken together, the analyses of both the main set and the core set demonstrate a highly consistent and biologically coherent enrichment pattern centred on synaptic regulation and neuronal signalling. Across GO (BP, CC, MF), KEGG, Reactome, and Human Phenotype databases, both gene sets converged on pathways related to synapse organization, signal transduction, cytoskeletal remodelling, vesicle-mediated transport, and activity-dependent plasticity. The recurrent identification of Rho GTPase signalling, actin cytoskeleton dynamics, calcium-dependent pathways, and long-term potentiation highlights structural and functional synaptic plasticity as a central mechanistic axis.

Importantly, enrichment of neurodegeneration-associated pathways and cognition-related human phenotypes further supports the disease relevance of these gene networks. While the main set reflects a broader synaptic regulatory architecture, the core set represents a more tightly defined subset reinforcing the same functional themes, thereby strengthening the robustness of the biological signal.

Collectively, these findings support a model in which RNA editing–associated genes are embedded within interconnected synaptic and signalling frameworks that regulate neuronal communication, structural stability, and cognitive function. The convergence across independent gene sets and annotation systems provides strong evidence that RNA editing alterations are positioned within molecular circuits directly relevant to synaptic dysfunction and neurodegenerative processes.

Comparison with genome-wide significant Alzheimer’s disease GWAS loci revealed no significant enrichment. The observed overlap did not exceed random expectation, suggesting that disease-associated RNA editing events in Alzheimer’s cortex may represent a regulatory mechanism largely independent of inherited genetic susceptibility loci.

## Supporting information

Supplemental Table-main set

Supplemental Table-core set

Supplemental Plot-main set

Supplemental Plot-core set

